# Light-Activated Cell Identification and Sorting (LACIS): A New Method to Identify and Select Edited Clones on a Microfluidic Device

**DOI:** 10.1101/204693

**Authors:** Annamaria Mocciaro, Theodore L. Roth, Hayley M. Bennett, Magali Soumillon, Abhik Shah, Joseph Hiatt, Kevin Chapman, Alexander Marson, Gregory Lavieu

## Abstract

CRISPR-Cas9 gene editing has revolutionized cell engineering and promises to open new doors in gene and cell therapies. Despite improvements in the CRISPR-editing molecular toolbox in cell lines and primary cells, identifying and purifying properly edited clones remains slow, laborious and low-yield. Here, we establish a new method that uses cell manipulation on a chip with Opto-Electronic Positioning (OEP) technology to enable clonal isolation and selection of edited cells. We focused on editing *CXCR4* in primary human T cells, a gene that encodes a co-receptor for HIV entry. T cells hold significant potential for cell-based therapy, but the gene-editing efficiency and expansion potential of these cells is limited. We describe here a method to obviate these limitations. Briefly, after electroporation of cells with *CXCR4*-targeting Cas9 ribonucleoproteins (RNPs), single T cells were isolated on a chip, where they proliferated over time into well-resolved colonies. Phenotypic consequences of genome editing could be rapidly assessed on-chip with cell-surface staining for CXCR4. Furthermore, independent of phenotype, individual colonies could be identified based on their specific genotype at the 5-10 cell stage. Each colony was split and sequentially exported for immediate on-target sequencing and validation, and further off-chip clonal expansion of the validated clones. We were able to assess single-clone editing efficiencies, including the rate of monoallelic and biallelic indels or precise nucleotide replacements. This new method will enable identification and selection of perfectly edited clones within 10 days from Cas9-RNP introduction in cells based on the phenotype and/or genotype.

## Introduction

Cell engineering through gene editing is fundamentally a two-step bioprocess: upstream, delivery of genome editing machinery to the cell type of interest to generate efficient and specific edits; and downstream, identification and selection of the cells that have been properly edited.

CRISPR-Cas9-mediated gene editing is a powerful tool to engineer cells lines and primary cells (1–3). The method enables precise correction or introduction of mutations within an endogenous genomic locus through co-delivery of a DNA template for homology-directed repair (HDR). There are widespread efforts to use this approach in clinically relevant systems to model genetic disorders (4) and for gene therapy to correct disease-driving mutations (5).

Many research and therapeutic applications are currently limited by the low efficiency of precise HDR-based editing. Even with improved delivery of Cas9, some targeted cells remain unedited. In addition, Cas9-mediated DNA breaks are repaired frequently by Non-Homologous End Joining (NHEJ) mechanisms that can introduce varying insertion and deletion mutations (indels) at the cut site resulting in undesirable editing outcomes (6, 7). Precise editing is complicated further because two copies of somatic alleles are present in the diploid genome. Therefore, in a given cell, HDR-mediated editing might occur only on one allele while the other allele is either unedited or imprecisely edited by NHEJ-mediated repair. Progress has been made to enhance the efficiency of HDR-based editing (8), however a technology to identify cells with desired monoallelic or biallelic edits is urgently needed to realize the full potential of CRISPR.

Selection of edited cell clones currently relies on limiting dilution or Fluorescence-Activated Cell Sorting (FACS)-based single-cell sorting to isolate single cells. When genome editing induces a phenotypic alteration that is detectable by fluorescence (i.e. cell surface expression of a target that can be non-lethally assessed with fluorescently-labeled antibody), FACS provides a method of enriching edited cells (9), significantly narrowing the number of clones to propagate and analyze. However, when the desired edit is phenotypically silent, a larger number of clones need to be isolated for subsequent sequencing to ensure that at least one of them has been properly edited. Moreover, even though high-purity cell sorting can be achieved, viability after sorting is often low to moderate, especially for cell types that are particularly sensitive to hydrodynamic stress or low-density culture conditions (e.g. primary cells or pluripotent stem cell lines). As a consequence, investigators often need to isolate a large number of clones and then proceed with tedious and time-consuming efforts to expand all of them individually. Each clonal line must then be assessed by sequencing to find those that bear the desired edits. Generating validated clonal lines can require several weeks. Therefore the development of a method that allows screening of edited cells and minimizes cell manipulation and hands-on culturing would constitute a significant addition to the current genome engineering toolbox.

Here we present proof-of-concept data highlighting the abilities of a new platform that integrates mechanical, fluidic, electrical and optical modules to enable single-cell manipulation, clonal expansion and phenotypic analysis in nanoliter volumes. The platform takes advantage of the Opto-Electronic Positioning (OEP) technology, which allows light-controlled manipulation of single cells (10–12). OEP is based on the principle of light-induced dielectrophoresis (DEP) force, an electrical gradient force. The OEP-microfluidic device (the “chip”) consists of a transparent electrode on a silicon substrate with a fluidic chamber sandwiched between the two. The substrate is fabricated with an array of photosensitive transistors. When focused light hits the transistors and a voltage is applied, a non-uniform electric field is generated. This imparts a negative DEP force that repels particles (including cells) using light-induced OEP (Fig 1 A). In the absence of targeted light, no force is generated; when light is shined on the photoconductive material, DEP force is generated and cells trapped inside light “cages” can be moved across the chamber. In addition, nano-pens are integrated into the chip to isolate cells from each other, enabling on-chip culture of well-separated colonies emanating from single cells (Fig 1 A and material section for more details).

**Figure 1:**
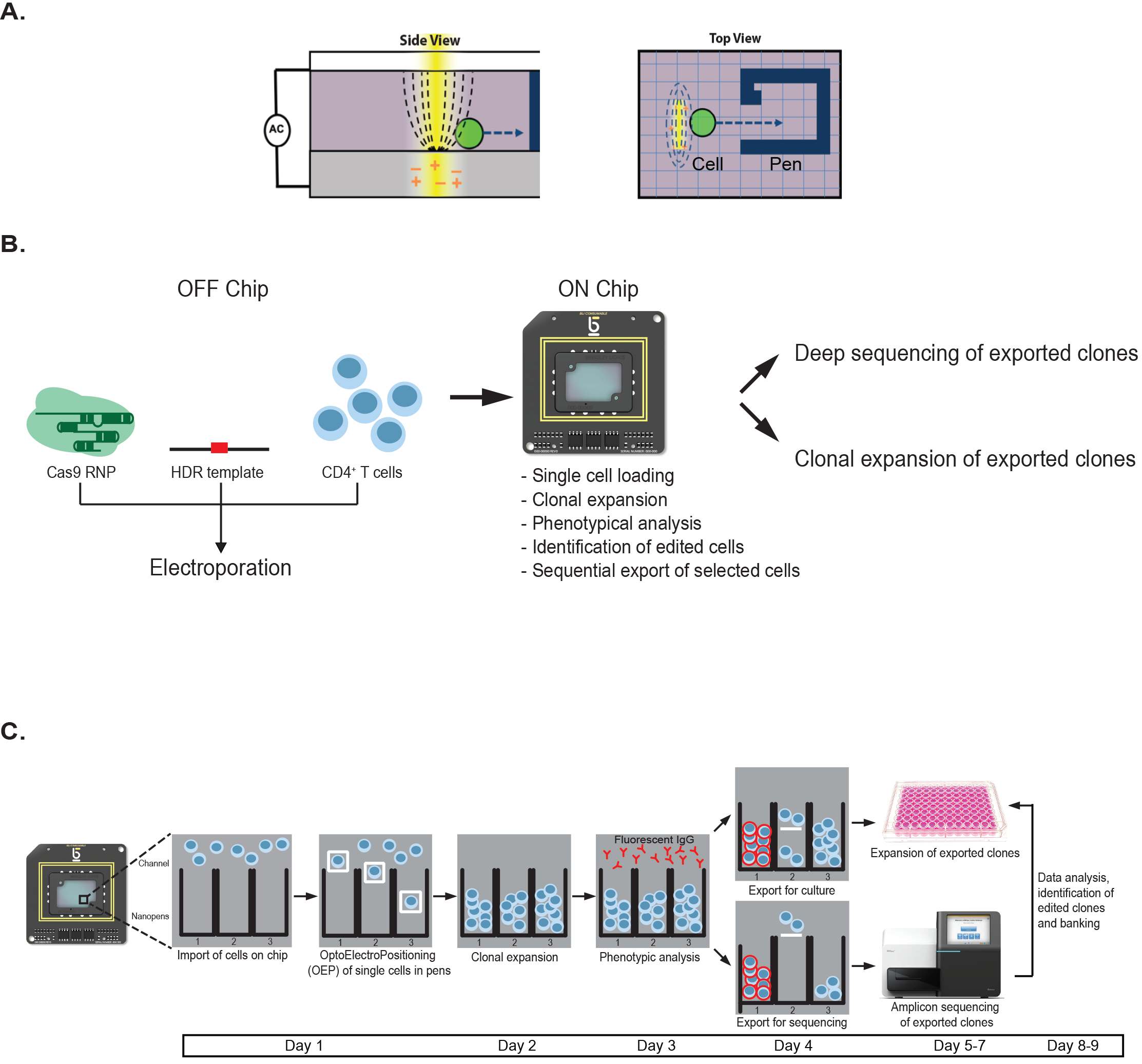
Method to identify and select edited cell with high precision. A) Schematic. Side (left panel) and top (right panel) views of the chip, depicting the OEP principle. A single cell (green) is moved inside a nano-pen (blue lines) through OEP (white arrow). B and C) Schematic representation of the LACIS workflow. T cell electroporation is performed off-chip, while clonal expansion, phenotype assessment, and export are performed on-chip. Each colony is split and exported. The first half of the colony is exported and further expanded through off-chip culture, while the remaining half is exported for validation through amplicon sequencing of the *CXCR4* locus. After on target validation, the desired clones are selected for further expansion and banking.

The advantages of the OEP-integrated platform include the capacity for: 1) massive parallel cell manipulation; 2) on-chip clonal expansion through absolute control of CO_2_, temperature and media perfusion; 3) on-chip fluorescence-based phenotypic assessment; and 4) sequential export of clones of interest for downstream processing (Fig 1 C). Every step of the workflow is defined and the process is highly automated such that it can be operated in a >90% hands-off manner. This new platform has allowed us to develop a method that facilitates both identification and selection of properly edited cells, including human primary T cells, as shown in the experiments presented below.

Here, we interrogated individual T-cell colonies on-chip after electroporation. Up to 50% of single T cells loaded on chip proliferated into a colony and fewer than 20% of the cells electroporated with *CXCR4* editing reagents had detectable CXCR4 cell surface labeling (vs. 80-90% CXCR4+ in control T cells electroporated with scrambled gRNA). After export of selected clones from the chip, further genotypic assessment through on-target sequencing revealed that approximately 5% of the putative edited candidates had bi-allelic HDR-based edits, and more than 50% of the exported clones were able to proliferate. The proposed method enabled the identification and the final selection of those precisely edited clones.

## Results

### Transfection, on-chip clonal expansion, and phenotype assessment

As previously described, human primary T cells were transfected with Cas9 ribonucleproteins (RNPs) targeting *CXCR4*, a gene encoding a surface receptor that acts as a coreceptor for HIV (9). The RNP complex was mixed with a short ssDNA oligonucleotide HDR template designed to replace 12 nucleotides within *CXCR4* (Fig 1 B) and impair cell surface expression. We previously reported up to ~20% HDR efficiency at this locus (9) based on deep sequencing analysis of a bulk population of edited cells. However, bulk sequencing of alleles from a cell population cannot distinguish the portion of mono-and bi-allelic knock-ins at the single-cell level. To obtain both phenotypic and genotypic data from individual edited clones, T cells were imported onto the chip one (Day 1) or four days (Day 4) after electroporation with CXCR4 Cas9 RNPs (Fig 2 A). We assessed editing efficiency at these two time points to identify further timeline compression options. After loading, flow was stopped to keep cells immobile within the main channel, which distributes media to multiple nano-pens (up to 3500/chip) through diffusion. Single cells were automatically selected and trapped into light cages that enable single cell positioning within the nanopens, in 17 out of the 18 fields of view (FOVs) that are visualized on the chip (Fig. 2 A and E). Non-penned cells remaining within the channel were flushed out of the chip. Importantly, we performed a second import with T cells electroporated with RNPs containing a scrambled control gRNA that does not target any locus in the human genome, positioning them in the remaining FOV (Fig 2 E). After three days of culture, during which fresh media was perfused into the main channel, we assessed on-chip clonal expansion. We first identified the pens that were initially loaded with single cells (to ensure clonality), and counted the number of pens that contained >6 cells after 3 days of culture. We established that, across multiple chips, approximately 15% or 40% of single cells, loaded at day 1 or 4, respectively, formed a colony (Fig 2B). The size of the individual colonies was heterogeneous (Fig 2 E, blue circle). The average doubling time was about 18 hrs over 3 days of growth with no significant delay in cell division timing (data not shown). These data strongly suggest that manipulation by OEP does not impair cell viability, and that diffusion of nutrients from the channel to the nano-pens maintains cell growth at expected levels. Importantly, we used nano-pens that were initially empty to track putative on-chip cross-contamination (cell transferred from one pen to another). Fewer than 2% of initially empty pens acquired cells within the three days of culture, indicating greater than 98% on-chip clonality (Fig S1). This rare cross-contamination that was observed might be explained by the high motility of activated T cells.

**Figure 2:**
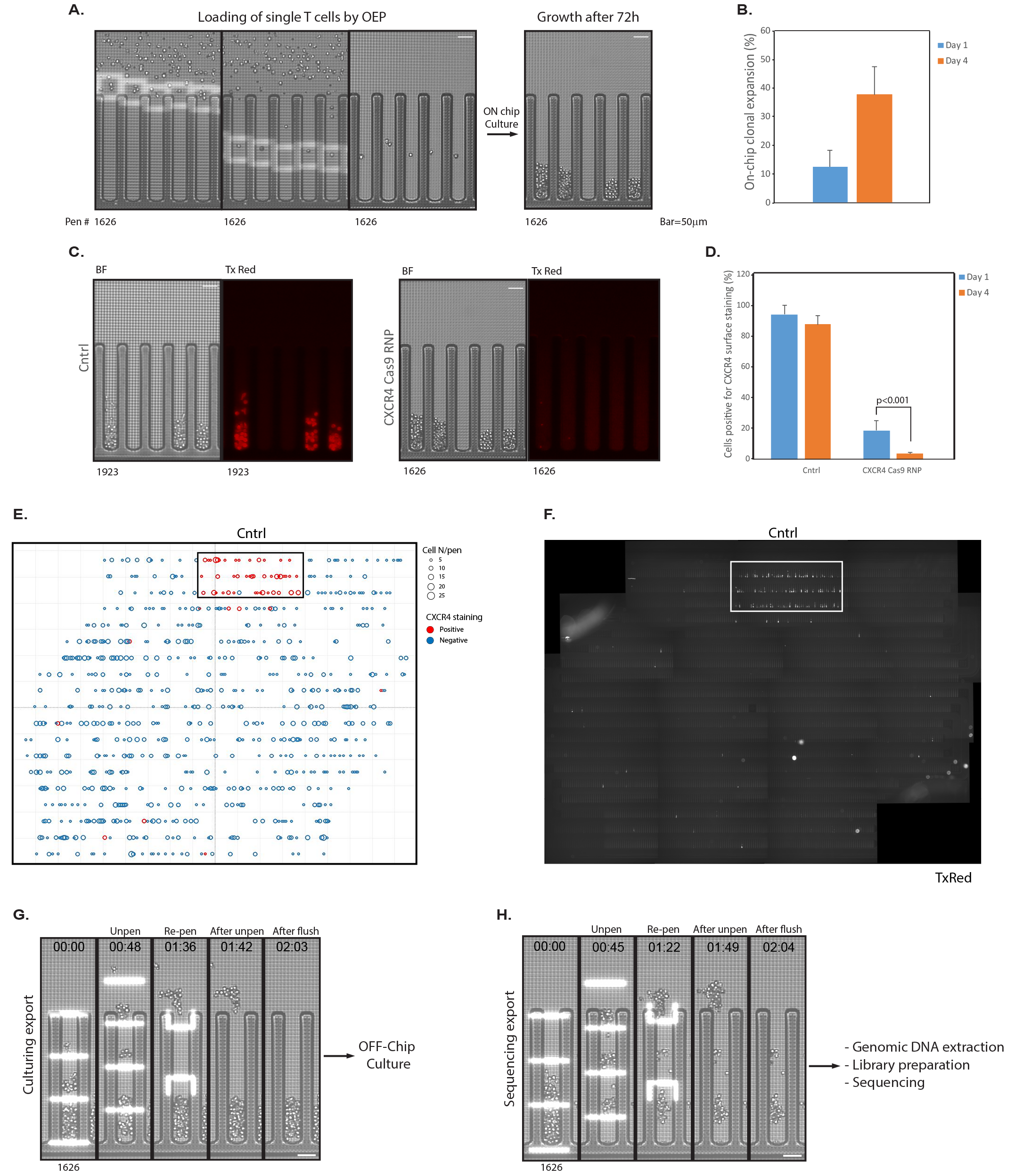
On-chip clone expansion, identification and selection. A) Electroporated cells are loaded on chip. Single cells located in the channel are automatically identified and captured within light cages and positioned into nano-pens, then cultured for 3 days. B) On-chip clonal expansion of clones loaded 1 or 4 days after electroporation. Results shown are mean±SD of three independent experiments (N≥9 chips, >300 clones analyzed per chip). C) On-chip phenotype assessment with fluorescently-labeled anti-CXCR4 antibody. Left panel, control cells. Right panel, putative edited candidates, negative for CXCR4 surface staining. D) CXCR4 staining for control cells and putative edited candidates loaded 1 or 4 days after transfection. Results shown are mean±SD of three independent experiments using primary T cells isolated from 4 different donors (N≥6 chips, >300 clones analyzed per chip). E) Graphic representation of on-chip clonal expansion and CXCR4 staining as a function of on-chip positioning. Each circle represents a single colony within a nano-pen, automatically identified by X,Y coordinates. The diameter of the circle is proportional to the colony size. Clones positive for CXCR4 are depicted as red circles. The heavy box indicates the FOV reserved for control, scrambled control RNP-treated cells. F) Composite image of the entire chip in the TxRed channel, showing CXCR4 staining. 18 fields of view were assembled together. White rectangle shows control cells transfected with scrambled control Cas9 RNPs and loaded within a single FOV. The 17 other FOVs contain clones electroporated with CXCR4 Cas9 RNPs. G-H) Representative images of the split export process: during Culture Export, the first half of the colony is unpenned, pushed into the channel, and exported into a well of a 96-well plate. For sequencing export, the second half of the colony is exported into a second 96-well plate for next generation sequencing and on-target validation.

Next, we established an on-chip phenotypic assay to identify clones that had undergone successful *CXCR4* editing. Fluorescently-labeled anti-CXCR4 antibody was imported into the chip, and media flow was interrupted to allow diffusion of the antibody into the pens. After 45 min of incubation, the chip was continuously flushed for 30 min with fresh media, to remove excess free antibody. Fluorescent images of the entire chip were taken (Fig 2 C and F) and the number of colonies positive for CXCR4 surface expression was quantified (Fig 2 D). Among the colonies formed by control cells across all chips, roughly 95% (day 1) and 85% (day 4) of clones were positive for CXCR4 (Fig 2 C, D). Strikingly, for CXCR4-edited cells loaded 1 day after electroporation, only 20% of the colonies showed presence of CXCR4 on the cell surface. In cells from healthy donors loaded 4 days post-electroporation, the number of colonies positive for CXCR4 staining dropped to around 5%. Importantly, each single pen was assessed for colony formation and fluorescence signal and a report was automatically generated to identify the nano-pens containing the clones of interest (Fig 2 E and F).

### Split-Export, On-Target Validation and Selection

Among all the putative edited clones that were automatically identified we selected a short list of candidates to export for on-target validation through next generation sequencing (NGS; 48 clones exported per chip, 9 chips in total). Our goal was to validate as early as possible the desirable clones in order to avoid wasting hands-on culturing efforts on clones that were not properly edited. To achieve this, we developed a pipeline that enabled a “split export” for clones of interest. Briefly, for each selected colony, roughly half of the cells were moved from the nano-pen into the channel via light bars (Fig 2 G). Un-penned cells (>5 cells/colony) were flushed out and collected in a defined well of a 96-well plate kept in a CO_2_- and temperature-controlled incubator for further off-chip culture. We termed this step “culture export.” Cells were exported from 48 nano-pens of each chip in this manner. We inserted 48 control blank exports (from empty nano-pens) between each clonal export to assess cross-contamination between wells introduced during and after export. Following culture export, media was replaced with Export Buffer and remaining cells from each nano-pen’s colony were serially transferred in the main channel and flushed out within a small volume of buffer into a corresponding well of a 96-well PCR plate kept at 4□C. We termed this step “sequencing export.” Efficiency of the export process, defined as the fraction of nanopens from which more than 1 cell was transferred to the channel, was **greater than** 80% (Fig S1 B).

Immediately after the sequencing export, collected cells (>5 cells per colony) were lysed and prepared for deep sequencing of the *CXCR4* locus (Fig S2 A and B). The sequencing reads from each individual clone were then aligned to the *CXCR4* WT sequence (Blue), the predicted HDR sequence (Green), or neither (called as a NHEJ due to introduced indel or point mutations, Orange) (Fig S2 C). Aggregating all the alleles found in cells from clones isolated on-chip on either day one or day four post electroporation allowed for a genotype to be assigned to each clone (Fig 3 A and B). In one healthy human blood donor, clones could be identified that possessed a variety of genotypes, from no edits at all (WT/WT), to mixed alleles of NHEJ-introduced indels, to mono-allelic HDR (with either WT sequence or indels on the other allele), to bi-allelic HDR (HDR/HDR) (Fig 3 A and B). Of note, not all *CXCR4* edited clones identified with loss of CXCR4 surface expression had 100% editing at the targeted CXCR4 locus, potentially due to Cas9 steric hindering CXCR4 transcription but not inducing a noticeable cut, large deletions unable to be identified by amplicon sequencing, or other unknown factors. More than two individual alleles were found in some clones, potentially due to editing events occurring after the first cell division (i.e. four alleles now present that could be edited), or cross-contamination between wells during culture, export, or NGS library preparation (Fig S3).

**Figure 3:**
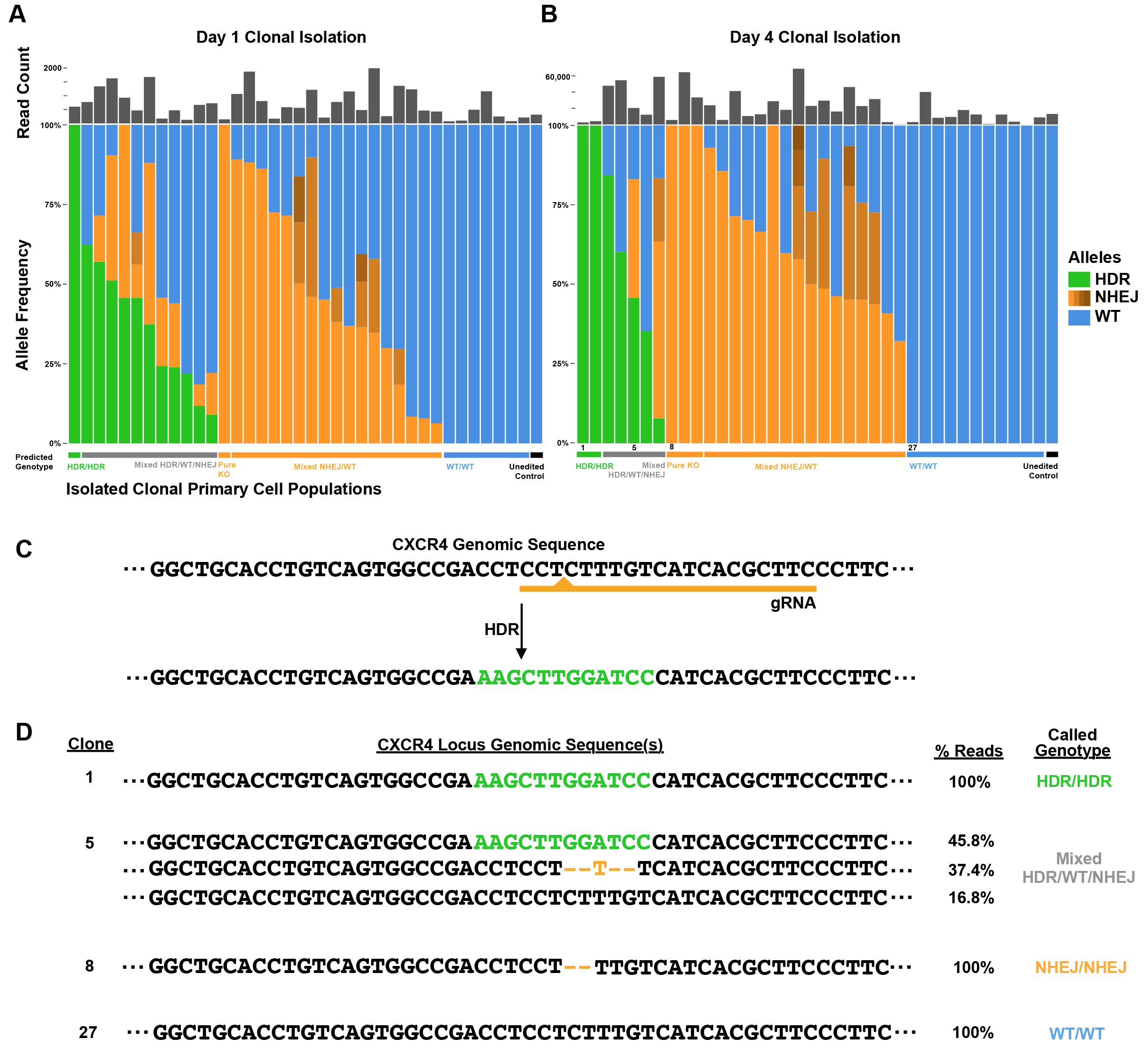
Off-chip sequencing of editing outcomes in individual clones. A-B) Proportions of reads mapping to HDR (Green), NHEJ (Orange), or WT (Blue) editing outcomes in each individual clone isolated and sequenced from cells loaded on-chip either 1 day (A) post-electroporation or 4 days (B) post-electroporation. The total read count from each clone in the sequencing run is displayed above the allele frequency. CD) Clones with many different genotypes can be identified, including those that integrated the HDR template (C) on both alleles (100% HDR, Clone 1), as well as clones with the same NHEJ edit on both alleles (Clone 8 with a two base-pair deletion), or mixed genotypes with more than two alleles present (Clone 5, potentially due to CRISPR editing continuing to happen after an initial round of division post single cell isolation).

Sequencing a portion of a clonal population while maintaining an ongoing culture of cells from the same colony allowed for clones to be identified based on their genotype, such as bi-allelic HDR (Fig 3 C). Selected examples of genotypes of clones isolated day four post-electroporation demonstrate the ability to identify such bi-allelic HDR integrations (Fig 3 D). To assess the fidelity of the off-chip sequencing and confirm the short ssDNA HDR template was not causing sequencing artifacts, we sequenced several individual unedited control clones (Black, unedited controls electroporated with a scrambled gRNA-based Cas9 RNP as well as the same HDR template as used for CXCR4-edited cells) that had been loaded in a pre-determined area of the chip and exported (Fig 2 E and F). As expected, greater than 97% of control clones showed no genomic alteration in the targeted *CXCR4* locus (WT/WT genotype, Fig 3 A and B). Overall, sequencing revealed that bi-allelically edited HDR clones could be identified while maintaining a live culture of the same clones.

Independently, we then assessed the post-export viability within the “Culture Export” plate. Exported clones were maintained for an additional week in culture, then plates were imaged and colony formation was quantified (Fig S4 A and B). Depending on the export conditions, up to 80% of the exported clones were able to survive and expand, with an average **of roughly** 60% of viability across all chips. Notably, we observed some variability in colony survival rates after export. In one case the off-chip post export viability was below 10% (data not shown). In that particular case, the number of cells exported from each pen was on average less than 5. We then refined our analysis, and we observed a strong correlation between the off-chip colony survival rate and the number of cells exported from each nano-pen (Fig S4 B). We concluded that, with current protocols, at least 10 cells needed to be exported for further off-chip clonal expansion in order to ensure greater than 50% post-export viability.

With approximately 5% bi-allelic HDR editing at the *CXCR4* locus and greater than 50% post-export viability, our results suggest that as few as 100 clones could be screened for on-target sequencing validation to ensure that at least 1-2 precisely edited primary human T cell clones are collected after culture export and will survive clonal expansion. This method is immediately relevant to identify and bank accurately edited clones of human primary cells.

## Discussion

Here we demonstrated that the Light-Activated Cell Identification and Sorting (LACIS) method is well suited to isolate clones that have been properly edited with precision. Compared to other methods, LACIS provides multiple advantages: this workflow removes the wasteful hands-on cell culture effort on undesired clones that are not properly edited. In addition, desired clones are identified quickly (<10 days), allowing for increased iterations and faster bioprocess optimization. Exporting larger numbers of cells per clone directly improves viability and expansion of the selected clones, and therefore contributes to increase the overall process efficiency (Fig S4). Importantly, this workflow can be almost fully automated which will enable significantly enhanced scale relative to current protocols.

In this study, we focused on primary human T-cell editing. We showed that the current capacity of the chip enables the identification of bi-allelically HDR-edited T cells, which at the targeted *CXCR4* locus was approximately 5% of edited cells. Therefore, even for a low-efficiency edit, the presented workflow is advantageous and should guarantee successful selection of cells with the desired genotype, whether or not edited cells can be phenotypically selected.

The present study is the first demonstration of a broadly applicable method that will enable selection of edited cells based on genotype and/or phenotype. The initial use of FACS enabled only a modest 4-fold enrichment of a certain cell sub-type based on one fluorescent criteria (13), but now - nearly 50 years later - enrichment can reach thousands of fold and allows multi-parametric analysis of heterogenous cell populations. This offers some perspective for future improvements in experimental throughput that will require innovative design of the chip to enable massive parallel genotyping and phenotyping throughout the entire chip (>1000 clones) within each run.

In our study, we primarily focused on the genotypic validation of edited clones through targeted DNA sequencing. However, further development of our platform would enable to analyze mutation-induced perturbations at the whole-transcriptome level through RNA sequencing, thus introducing an additional way of linking genotype and phenotype, which is critical to understand disease genetics and characterize new therapeutic targets.

Recent improvements in NGS - especially barcoding and low input (5-20 cells) processing - are driving cost reductions that will enable larger scale characterization of edited cells while also assessing off-target effects when necessary. This promises to greatly facilitate and improve the manufacturing of edited cell lines for the scientific and medical community. In addition, our flexible platform could enhance other gene-editing workflows. For instance, our pipeline could facilitate the study of genetic disorders through the generation of heterozygous or homozygous model cell lines (ESCs or iPSCs) bearing knock-in disease mutation (17–19).

Much remains to be done to truly revolutionize the development of edited cell clones. For instance, culture and export of adherent cell lines need to be enabled, since they constitute many relevant models for disease. In addition, gene and cell therapy currently require dealing with a very large number of cells, which will require design improvements to this platform. Automation, new microfluidic layouts and integration of relevant technologies will enable high-throughput sorting of cells based on genotype with absolute precision.

## Materials and methods

### Human T cell isolation and culture

Primary human T Cell culture and RNP editing has been previously described (7). Briefly, PBMCs were isolated using SepMate tubes (STEMCELL) per manufacturer’s instructions from blood from healthy human donors under a UCSF CRB approved protocol. CD3+ T cells were negatively isolated from PBMCs using an EasySep (STEMCELL) negative magnetic isolation ket per manufacturer’s protocol. T cells were stimulated with plate bound CD3 (10 ug/mL, Tonbo Biosciences, clone UCHT1) and soluable CD28 (5 ug/mL, Tonbo Biosciences, clone CD28.2) antibodies at 1 million cells per 1 mL of RPMI media with 10% FBS. After electroporation, T cells were stimulated with CD3/CD28 dynabeads (Cell Therapy Systems, 1:1 bead to cell ratio) and 20 U/mL of IL-2 (UCSF Pharmacy) again at 1 million cells per mL of media until import onto the Optoselect chip.

### Cas9 RNPs electroporation

A two-component gRNA system was used-crRNAs targeting either *CXCR4* (target sequence 5’ to 3’: GAAGCGTGATGACAAAGAGG) or no human genomic sequence (“Scrambled” gRNA, 5’ to 3’: GGTTCTTGACTACCGTAATT) were synthesized (Dharmacon) and resuspended in 10 mM Tris HCl pH 7.4 with 150 mM KCl to a final concentration of 160 uM. tracrRNA was similarly synthesized and resuspended. The crRNA and tracrRNA were mixed 1:1 by volume and incubated for 30 minutes at 37C to produce 80 uM gRNA. 40 uM SpCas9 (QB3 Macrolab) was added at 1:1 by volume to the gRNA (a 1:2 molar ratio of Cas9 to gRNA) and incubated for 15 minutes at 37C to yield a 20 uM RNP. RNPs were prepared immediately before electroporation into T cells. A short ssDNA HDR template (ssODN) to insert a defined 12 bp sequence into *CXCR4* was chemically synthesized (IDT) and resuspended in nuclease-free H2O at 100 uM. The same *CXCR4* targeting HDR template (DNA sequence 5’ to 3’: GGG CAA TGG ATT GGT CAT CCT GGT CAT GGG TTA CCA GAA GAA ACT GAG AAG CAT GAC GGA CAA GTA CAG GCT GCA CCT GTC AGT GGC CGA AAG CTT GGA TCC CAT CAC GCT TCC CTT CTG GGC AGT TGA TGC CGT GGC AAA CTG GTA CTT TGG GAA CTT CCT ATG CAA GGC AGT CCA TGT CAT CTA CAC AGT) was used for both *CXCR4* and Scrambled gRNAs. Two days following stimulation, T cells were harvested and resuspended in P3 electroporation buffer (Lonza) at a concentration of 1 million cells per 20 uLs of buffer. 5 uLs of RNP (100 pmols) were added to 20 uLs of cells (1 million T cells) along with 1 uL of HDR template (100 pmols) were mixed and electroporated in a single well of a lonza 4D nucleofection system cuvette using program EH-115. Immediately following electroporation, 80 uLs of pre-warmed culture media were added directly to the cuvette and the cells were allowed to rest for 15 minutes in a 5% CO2 37C incubator for 15 minutes in the cuvettes before being stimulated and transferred out for further culture (see Human T Cell Isolation and Culture).

### Preparation of cell suspension for penning in Optoselect chip

T cells were cultured for 1 day or 4 days after electroporation in culture media Media [RPMI-1640 (Gibco) supplemented with 2mmol/L Glutamax (Gibco), 10% (vol/vol) FBS (Seradigm), 2% Human AB serum (ZenBio) and 50IU/ml IL-2 (R&D Systems), in the presence of Anti-CD3/CD28 Dynabeads (Gibco)]. Prior to loading onto the chip, cells were resuspended in culture media supplemented with 10ng/ml IL-7 and IL-15 (PeProTech) at a final density of 5e6 cells/ml.

### Conditions for automated cell penning

Experiments were conducted on commercialized BerkeleyLights platforms and chips. After priming, chips were washed twice with de-ionized water and flushed 6 times with culture media. Cells were imported onto the chips and loaded as single cells into Nanopens using OEP with the following parameters - nominal voltage: 4.5 V; frequency: 1000 kHz; cage shape: square; cage speed: 8 μm/s; cage line width: 10 μm. Loading temperature was set to 36^0^C. Brightfield images of each chip were acquired automatically at the end of the loading process and a BLI proprietary algorithm was used to detect and count cells by pen.

### Culturing conditions and cell expansion quantification

Chips were maintained at a temperature of 36^0^C during culture. CO_2_-buffered culture media was perfused through the chip at a flow rate of 0.01 μl/sec. For primary cell growth assessment and automated counting, Brightfield images of the chips were taken at distinct time points to quantify On-Chip Clonal Expansion (OCCE), defined as the percentage of nanopens containing a single cell that grew into a colony of 6 or more cells after 72h of culture. Cross-contamination across each chip was determined as the percentage of initially empty pens that acquired cells during culture.

### On-chip T cell staining

Cell surface staining was performed with αCXCR4-PE (12G5; BioLegend). The antibody was imported into the chip at 1:250 dilution in culture media and incubated for 45 min at 36^0^C. After staining, chips were perfused for 30 min with culture media media, to remove the excess antibody, and then images were acquired in Brightfield (25ms) and Texas Red (1000ms) channels.

### Split export of edited clones

Three to four days after loading, clones containing >10 cells that showed negative staining for CXCR4 were sequentially exported for off-chip culturing and genotyping. 48 clones and 48 blanks were exported per chip. In the first step of the split export (culturing export), roughly half of each clone (5-20 cells) was transferred from the NanoPen to the channel using light bars generated by OEP, with the following parameters - nominal voltage: 4.5 V; frequency: 1000 kHz; bar speed: 5 μm/s; bar line width: 10 μm. Export temperature was set to 36^0^C, export was performed in culture media and cells were flushed in a 20ul package volume into a barcoded round-bottom, tissue culture treated 96-well plate containing 100 ul of culture media supplemented with 10ng/ml IL-7 and IL-15 per well. The plate was kept in an incubator at 36^0^C and 5% CO_2_ for the entire duration of the export. For the second step of the split export (genotyping export), culture media was replaced with Export Buffer [PBS (Gibco), 5mg/ml BSA (Fisher Scientific), 0.1% Pluronic F-127 (Life Tech)] by flushing the chip 10 times before starting the export. Then, the remaining cells from the previously exported pens were transferred to the channel by OEP with the following parameters - nominal voltage: 5 V; frequency: 1000 kHz; bar speed: 5 μm/s; bar line width: 10 μm. Export temperature was set to 36^0^C, export was performed in Export Buffer and cells were flushed in a 5ul package volume into a barcoded 96-well PCR plate (Eppendorf) containing 20ul of mineral oil (Sigma-Aldrich) and 5ul of Proteinase K buffer [(10 mM Tris-HCl pH 8, 0.1 M NaCl, 1 mM EDTA, 200 μg/ml proteinase K (Ambion AM2546)] per well. The PCR plate was maintained at 4^0^C for the entire duration of the export.

### Sample processing for next-generation sequencing

Genomic DNA was extracted from exported clones by incubating in Proteinase K buffer (0.1 M NaCl, 10 mM Tris HCl pH 8.0, 1 mM EDTA) for 30 min at 55□C, then for 20 min at 80□C to inactivate Proteinase K. The genomic region around the CRISPR/Cas9 target site for CXCR4 gene was amplified by PCR with primers positioned outside of the HDR repair template sequence (positioned to avoid amplification of exogenous template) for 10 cycles using KAPA HiFi Hotstart ReadyMix (Kapa Biosystems, KR0370) according to the manufacturer’s protocol (PCR primers listed in Supplementary Table 1). Primers contained inline sample-specific barcodes. Barcoded samples from each plate were pooled to concentrate and remove mineral oil using Zymo DNA Clean and Concentrator Column (Zymo research, D4004). Excess PCR primers were removed by incubating with Exonuclease I (NEB, M0293S) in 1X Exonuclease Reaction Buffer (NEB, B0293S) for 1h at 37□C, followed by enzyme inactivation for 20min at 80□C. Amplicon pools were reamplified by PCR for 15 cycles using a universal primer to add the sequencing adaptor and secondary barcodes to allow parallel sequencing of multiple amplicon pools. PCR products of the expected size were isolated with Select-A-Size DNA Clean and Concentrator (Zymo research, D4080) as sequencing libraries. Pooled barcoded libraries were sequenced with 300 bp paired-end reads on a MiSeq (Illumina) instrument using the 300 cycles v3 reagent kit (Illumina).

### Sequencing data analysis and HDR/indel identification

All computational and statistical analysis were performed using Python 2.7 and Unix-based software tools. Quality of paired-end sequencing reads (R1 and R2 fastq files) was assessed using FastQC (http://www.bioinformatics.babraham.ac.uk/projects/fastqc). Reads with sample-specific inline barcodes were demultiplexed using our home-brew python script for FASTQ files splitting. Reads were then mapped on both the wild type sequence and the expected HDR edited sequence of *CXCR4* using bwa version 0.7.15 (20) with default parameters. Alignments files were sorted and indexed using samtools version 1.3.1 (21, 22). Variants were called using freebayes version 1.0.2
(https://arxiv.org/abs/1207.3907), a Bayesian haplotype-based polymorphism discovery tool. Genotypes were determined for each colony based on the number of reads matching either the wild type sequence, the HDR sequence or containing variants to these two sequences with a quality above 30. Python scripts implementing the demultiplexing, alignment and genotyping are available from the authors upon request.

### BLI Platform and Chip overview

The OptoSelect™ platform takes advantage of the OptoElectroPositioning (OEP™) technology, which enables light-controlled cell manipulation. OEP is enabled by the generation of a dielectrophoretic force (DEP), which occurs when a polarizable particle is suspended in a non-uniform electric field.

The proprietary OptoSelect™ nanofluidic device consists of a top transparent electrode and a bottom silicon substrate with a fluidic chamber in between. The substrate is fabricated with an array of photosensitive transistors. When light shines on the transistors, and if voltage is applied, a non-uniform electric field is locally generated in the fluidic channel. This imparts a negative DEP force that repels particles (including cells) using light-induced OEP. In the absence of targeted light, no force is generated; when light is shined on the photoconductive material, DEP force is generated and particles trapped inside light “cages” can be moved across the chamber. The chip contains a main fluidic channel and 3500 individual NanoPens™ chambers, which hold a 0.5nL volume each. Media is perfused across the channel by a fluidic system, and it diffuses to the nanopens. Cells are loaded into the channel through an import needle, from a sample tube or from a well plate, and using light cages they are moved into the nanopens at a speed of 5-15mm/s (Fig 1 A). Single cells loaded into pens are isolated from each other, and perfusion of CO2-buffered media through the chip during culturing at 36^0^C enables the in-pen expansion of clones over time. The chip is placed on a 3-axis robotic stage and an upright microscope mounted on top of the stage allows image collection of the entire chip area at 4x or 10x magnification in both brightfield and fluorescent channels, to monitor cell growth, morphology and to perform phenotypical analyses. After characterization, selected clones can be exported off the chip for further processing. The export is the reverse of the import process, where desired cells are moved using OEP from single NanoPens into the main channel and flushed into a target well of a wellplate positioned inside a CO2- and temperature-controlled incubator.

The imaging system can provide both brightfield and fluorescent imaging. A 360nm LED source is used to illuminate the background and a 400 nm-700 nm white light lamp combined with a Digital micromirror device (DMD) is used to structure light in desired patterns for light actuated dielectrophoresis (DEP). The system uses an upright microscope with an automated lens changer to image at 4X and 10X magnifications and a linear cube slider to collect fluorescent images in wavelengths corresponding to Cy5, FITC and TxRED fluorescent channels. More information are available at https://www.berkeleylights.com/contact-us/.

## Footnotes

Conflict of Interest statements: An.M., H.B, M.S, A.S, K.C, and G.L are fulltime employees of BerkeleyLights, a private company. Al.M. serves as an advisor to Juno Therapeutics and PACT Therapeutics and the Marson lab has received sponsored research support from Juno Therapeutics and Epinomics.

An.M., T.R, H.B, J.H, performed experiments, An.M., T.R, H.B, M.S, J.H, Al.M and G.L, designed research work plan and experiments. An.M., T.R, H.S, M.S., Al.M and G.L analyzed data. All the authors wrote and edited the manuscript.

## Acknowledgements

We thank members of Marson lab. A.M. holds a Career Award for Medical Scientists from the Burroughs Wellcome Fund and is a Chan Zuckerberg Biohub Investigator. We relied on the Flow Cytometry Core at UCSF, supported by the Diabetes Research Center grant NIH P30 DK063720. T.L.R. and J.H. were supported by the UCSF Medical Scientist Training Program (T32GM007618). T.L.R. was supported by the UCSF Endocrinology Training Grant (T32 DK007418).

---

**Supplementary Figure 1:**
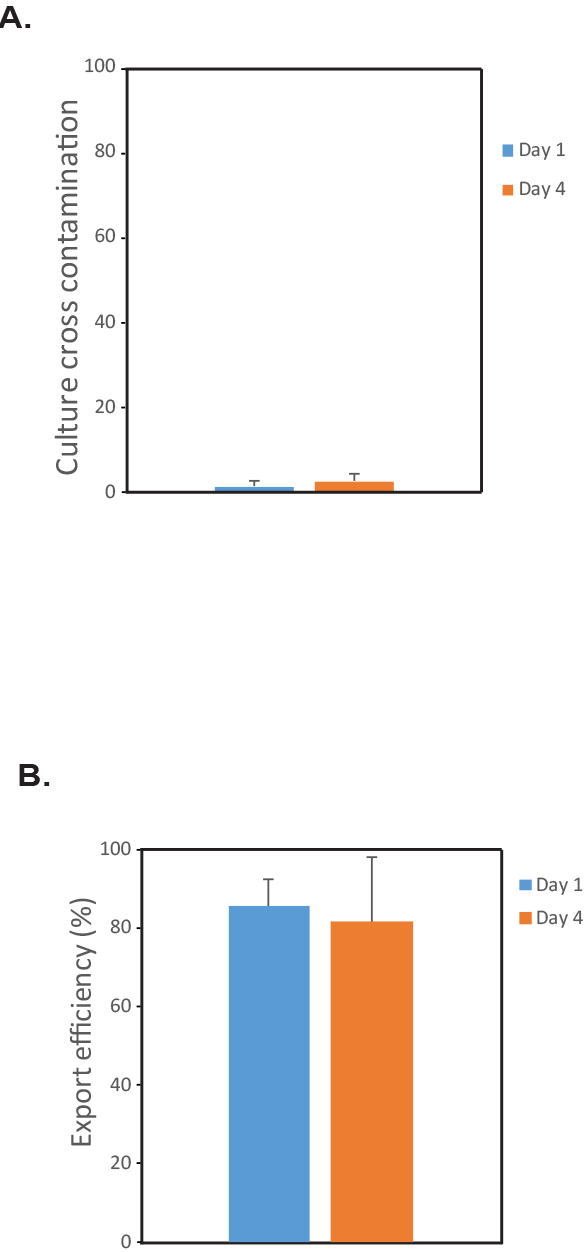
Assessment of cross-contamination and unpenning efficiency. A) Graph, Percentage of nano-pens that were originally empty and acquired unwanted cells over 3 days of on-chip culture. Results shown are means ±SD of three independent experiments (N≥6 chips, >100 NanoPens analyzed per chip). B) Unpenning efficiency for clones loaded 1 or 4 days after transfection. Results shown are means ±SD of three independent experiments (N≥6 chips, >20 clones exported per chip). Low level of cross-contamination observed may have occurred on-chip, during export or during preparation/sequencing of genomic amplicons.

**Supplementary Figure 2:**
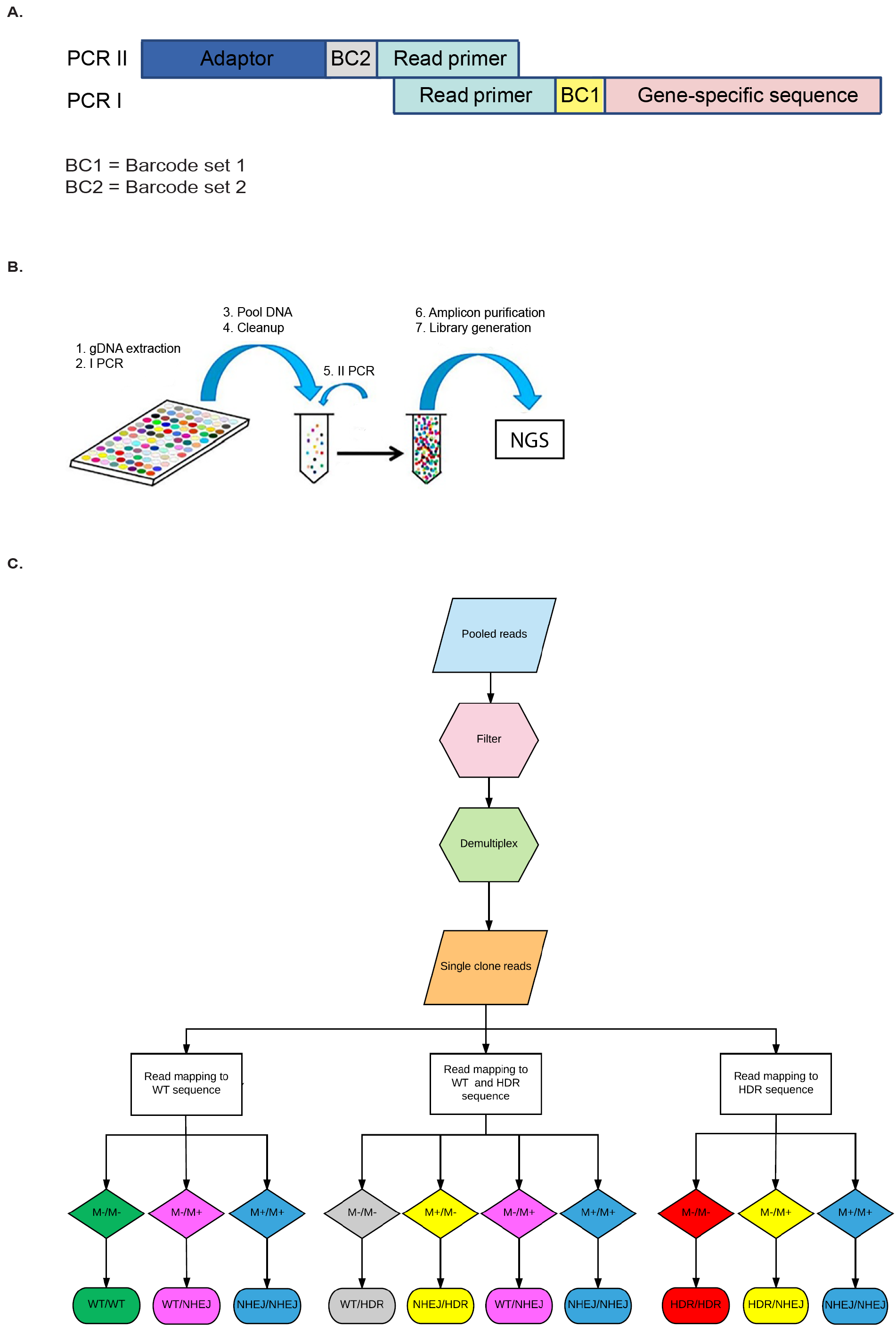
Sample processing for NGS and data analysis for clone genotyping. A) Organization of the sequencing primers used for NGS. The two-barcode system gives up to 96x96 index options using only two validated index sets. BC1=barcode set 1 (inline barcode, well identifier). BC2=barcode set 2 (plate identifier). B) Schematic representation of the molecular biology workflow for genomic DNA extraction and sequencing library preparation from the exported clones (See Materials and Methods). C) Flowchart of the NGS data analysis pipeline for genotype identification. After quality filtering and demultiplexing, reads from each clone were mapped to the reference WT or HDR sequence (M−). Clones presenting additional mismatches from the reference sequences (M+) in a 200bp region around the PAM site were identified as NHEJ.

**Supplementary Figure 3:**
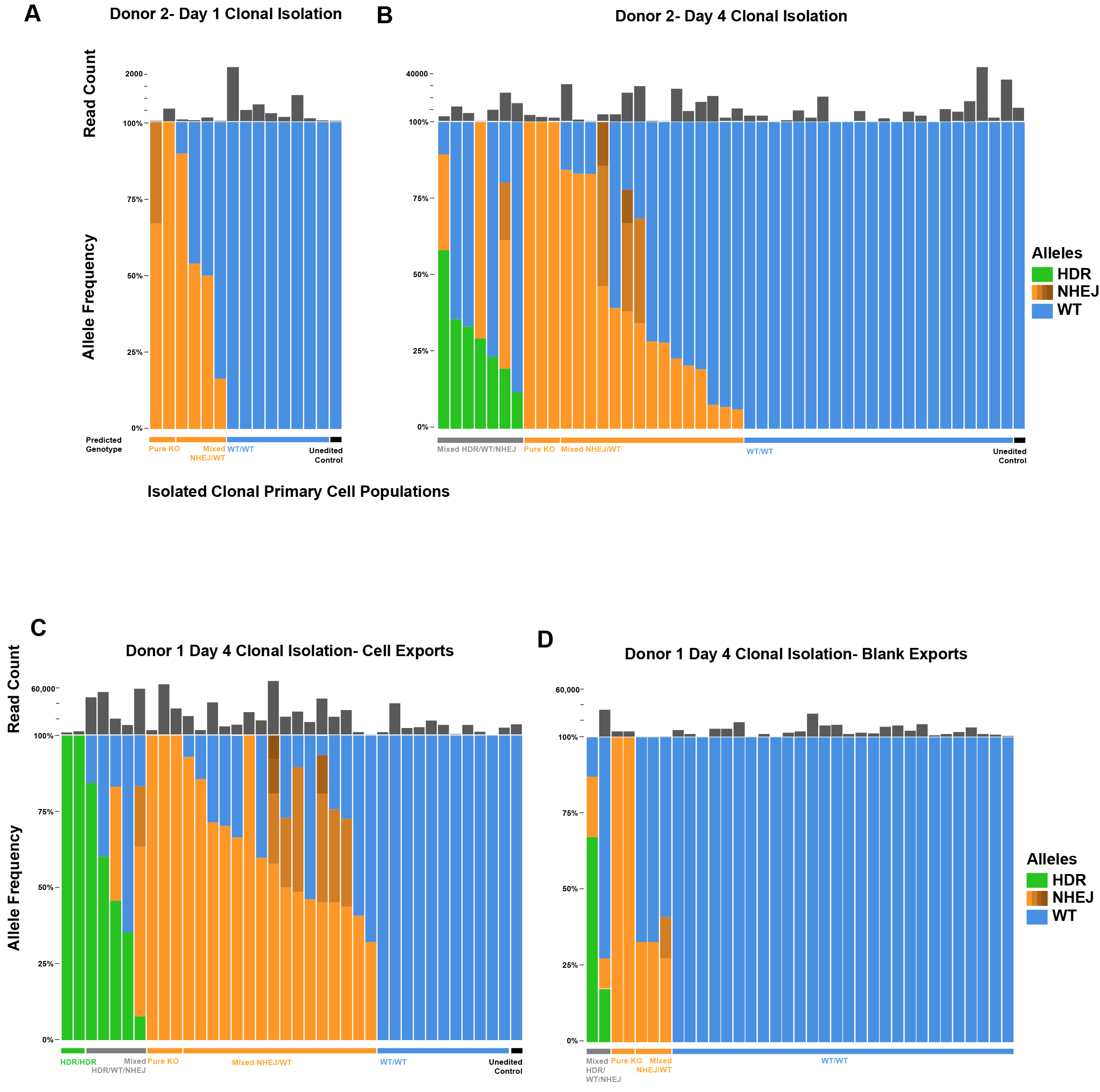
Off-chip sequencing of clones from second healthy human donor and export controls. A-B) Allele frequencies and read counts from cells isolated from a second healthy human blood donor and loaded on-chip either one day (A) or four days (B) after electroporation. C-D) Cells were exported off-chip for sequencing (C) but as a control for cross-contamination introduced during the export procedure media from on-chip wells that had no cells loaded (Blank controls, D) were also exported off-chip and processed similarly for NGS sequencing. The presence of reads in these blank wells indicates that some cross-contamination between wells occurred, either on-chip, during the export of cells off the chip, or during the genomic PCR amplication or NGS library preparation steps after export. This potential cross-contamination also could contribute to the detection of more than two different alleles in some wells of the cell exports (C)

**Supplementary Figure 4:**
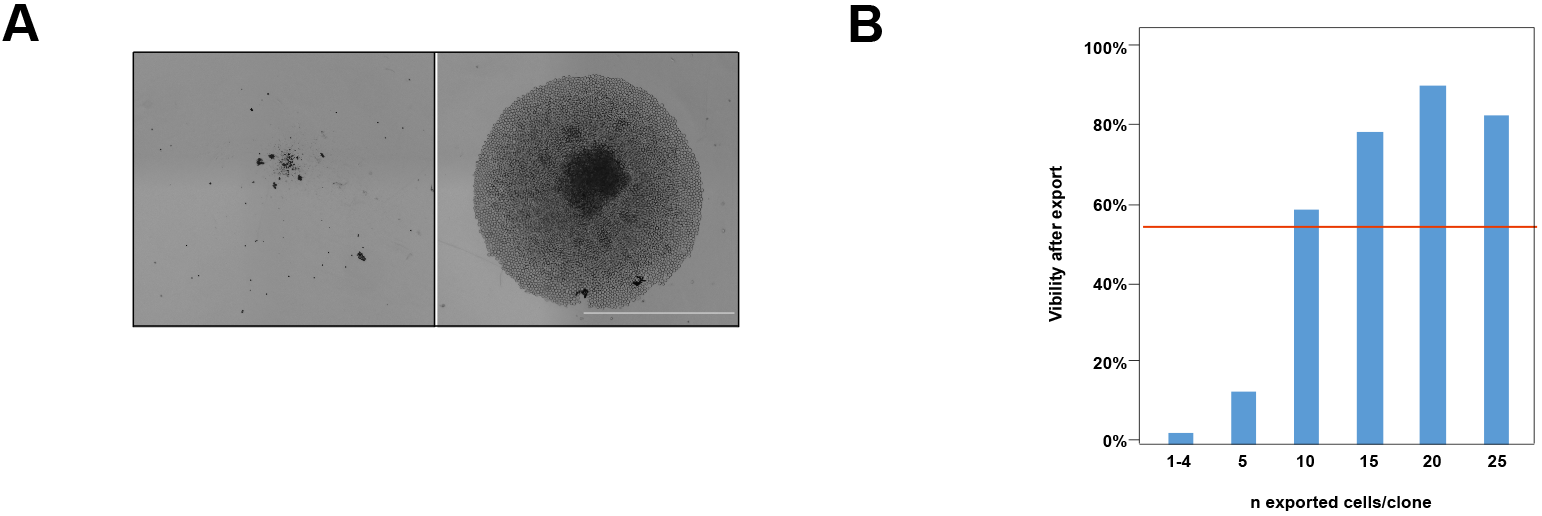
Off-chip clonal expansion of exported clones. A) Representative image of a colony after Culture export. Individual wells of the 96 well plates collected after export were imaged after 7 days in culture. Wells corresponding to blank exports were imaged as control B) Graph, cell viability (% of clones forming colony) after export as a function of the number of cells exported. The red line and the gray area indicate, respectively, the mean and the SEM of cell viability across the exported clones (N=363 clones analyzed).

